# Density-dependent private benefit leads to bacterial mutualism

**DOI:** 10.1101/2020.07.28.224550

**Authors:** Paul Jimenez, István Scheuring

## Abstract

Microorganisms produce materials leaked from the cell which are beneficial for themselves and their neighbors. We modeled the situation when cells can produce different costly secretions which increase the carrying capacity of the population. Strains that lose the function of producing one or more secretions avoid the cost of production and can exhaust the producers. However, secreting substances provides a private benefit for the producers in a density-dependent way. We developed a model to examine the outcome of the selection among different type of producer strains from the non-producer strain to the partial producers, to the full producer one. We were particularly interested in circumstances under which selection maintains partners that produce complementary secreted materials thus forming an interdependent mutualistic interaction.

We show that interdependent mutualism is selected under broad range of conditions if private benefit decreases with density. Selection frequently causes the coexistence of more and less generalist cooperative strains, thus cooperation and exploitation co-occur. Interdependent mutualism is evolved under more specific circumstances if private benefit increases with density and these general observations are valid in a well-mixed and in a structured deme model. We show that the applied population structure supports cooperation in general, which, depending on the level of private benefit and intensity of mixing helps either the specialist or the generalist cooperators.

## 1 INTRODUCTION

All animals live in close association with microorganisms. These microorganisms generally live in communities composed of many taxa and their genetically different strains. For example, the human colon microbiota is constituted by 100 to 300 different bacterial taxa together with dozens of fungi, archaea and viruses (Hillman et al., 2017). Also, different protists, bacteria, archaea, and viruses live in association with corals (Thompson et al., 2015; Bourne et al., 2016), whereas the bacteria covering the cuticle of two Neotropical ants belong to at least ten different phyla (Birer et al., 2017)

Besides the importance of microbes in hosts’ physical and mental health and even in their social behavior (Cho and Blaser, 2012; Mattoso et al., 2012; Valdes et al., 2018; Taylor, 2019; Sherwin et al., 2019), species rich soil microbiota are essential for maintaining the functioning of terrestrial ecosystems. They are involved in decomposing organic materials and in maintaining biogeochemical cycles of nitrogen and carbon (Bardgett et al., 2008; Jacoby et al., 2017).

One specific feature of microbial communities is that microbes release a great variety of leaky materials which can be enzymes (Gore et al., 2009), siderophores (Cordero et al., 2012), amino acids (Mee et al., 2014), antibiotic degrading molecules (Yurtsev et al., 2013) etc. The common nature of these extracellular substances is that they increase the producers’ fitness but are also beneficial for other cells in the vicinity. If producing the secretion is costly, then the cost of producing this material is borne only by the producers, while the benefit can be exploited by all cells in the neighborhood. Although the average fitness is higher in a population of producers than in a population of non-producers (cheaters), non-producers invade, supersede producers and finally become fixated in the population. Hence, leaky materials are public goods which generate social conflicts between microbes (Tarnita, 2017). Despite cheaters seeming to have a selective advantage over producers, numerous mechanisms (e.g. limited mixing and local interactions of individuals (Szabó and Hauert, 2002; Helbing et al., 2010), the nonlinear effect of public goods on fitness (Foster, 2004; Archetti and Scheuring, 2012, 2016), density dependence of public good (Rankin, 2007; Kokko and López-Sepulcre, 2007), the private benefit of producers (Drescher et al., 2014; Estrela et al., 2016) and information transfer among cells (Czárán and Hoekstra, 2009; van der Ploeg, 2005) can prevent it.

Most previous theoretical and experimental studies focused on situations when only one type of leaky material is secreted. However, numerous types of excreted substances can be distinguished in a complex microbiota. Thus, besides those producing all possible secretions and strains producing none of them, many other specialists (or partial) producers can be present simultaneously. If these specialist strains produce complementary materials, they form interdependent mutualistic networks. Although, there are studies on simple artificial communities with interdependent mutualist strains (Cavaliere et al., 2017; Mee et al., 2014; Harcombe et al., 2014; Pande et al., 2014) it is not known how frequently these mutualistic interactions appear in natural microbiota.

The situation is rather different if the produced leaky material is a costless waste product from some type of cells and is consumed by other cells and vice versa. This by-product mutualism is probably frequent among strains (Harcombe, 2010; Rivière et al.; Smith et al., 2019a), which can explain unculturability of most of the microbial species (Kaeberlein et al., 2002; Stewart, 2012). Some studies suggest that competitive interactions dominate over cooperative ones in microbial communities, i.e. mutualism is relatively rare in natural microbiota (Foster and Bell, 2012; Coyte et al., 2015). Here, we reveal circumstances under which costly interdependent mutualism is stable from ecological and evolutionary points of view in microbial communities.

Previously, Oliviera et al. (Oliveira et al., 2014) studied the evolutionary stability of interdependent mutualism in a model of microbial communities. In this paper, we revisit and extend their model, thus we present their assumptions and results in more detail. They considered a situation where cells can produce different leaky materials that are essential for a higher carrying capacity. Variants may range from non-producers (complete cheaters) to generalist strains producing all types of secretions. Oliveira et al. assumed that numerous local populations are founded initially by some cells. Populations grow within local habitats following a dynamics depending on the proportions of different producer strains. After the growth phase, cells are mixed together and establish new local populations which then start to grow. This life cycle continues iteratively until a stationary state is reached. They showed that specialist interdependent producer strains are selected only if the cost of secretion is high and a medium number of individuals found a new habitat (genotype mixing or heterogeneity is at intermediate level). A low level of heterogeneity favors generalist cooperators, while high levels of heterogeneity prefers complete cheaters. Additionally, the coexistence of complete cheaters and generalist cooperators was possible under at very specific circumstances in their model. Since the cost of producing leaky material is not high and a low level of heterogeneity can be found only in specific natural microbiota, the authors concluded that cooperation among generalists should be more common in nature than interdependent mutualism among specialists.

The dynamical model used by Oliveira et al. contains only one crucial parameter, the cost of producing secretions. This simplifies analysis and the interpretation of results, but naturally neglects important characteristics of the problem being examined. One of the most important simplifications is that the model does not consider the private benefit of the producer. Moreover, it neglects that this private benefit is density dependent.

Neglecting the producer’s private benefit assumes that the leaky materials are public goods available for all individuals equally, which is not necessarily true. In reality, there is inhomogeneity in the concentration of the secreted substances. Producer cells themselves and other cells in the producer’s neighborhood typically detect higher concentrations of secretions than cells farther away from the producers (Gore et al., 2009; Morris et al., 2014; Lindsay et al., 2018; Smith et al., 2019b).

Depending on the cell’s mobility and the diffusion rate of secretions, either high or low density is more beneficial for producer strains as compared to non-producer ones. If diffusion of the secretions is faster than the cells movement, a concentration gradient is present around the producers. Therefore, increasing the density increases the proximity of cheaters to the cooperators which helps cheaters to exhaust cooperators intensively (Ross-Gillespie et al., 2009; Morris et al., 2014). On the other hand, diffusion of the secretions can sometimes be slow compared to the cell’s movement. Then, increasing density enhances the amount of public goods that can be captured by the cooperators themselves (Dobay et al., 2014). Lindsay and his colleagues recently proved experimentally that both cases are realized (Lindsay et al., 2018). Thus, we extended the model used by Oliveira et al. (2014) to include density-dependent private benefit of cooperators. Besides this extension, we also modified the basic dynamical system to describe the competition among the strains more adequately.

We show that density-dependent private benefit of cooperation significantly widens those circumstances where selection prefers specialist cooperators over generalist cooperators and complete cheaters. Further, we show that specialist cooperators can typically coexist with generalists or complete cheaters, if the private benefit of cooperation decreases with density.

In the following, we introduce the dynamical model systems from the simplest model with one secretion to those involving any number of secretions, and then present the structured population model. The first part of the Results section contains the invasion analysis and numerical studies in the local populations. Then we investigate the dynamics of the structured model numerically. The Discussion interprets and summarizes the main results.

## 2 THE MODELS

### Population dynamics with a single secretion

First, we assumed that only one leaky material can be produced. There are two types of cells; type [0] which does not produce the leaky material and type [1] which does. The density of cheater (type [0]) and producer (type [1]) cells is denoted by *g*_0_ and *g*_1_, respectively. Cheaters replicate with a replication rate *r* and follow a logistic growth with carrying capacity *K*_*base*_. Thus the dynamics of cheaters without the presence of cooperators is

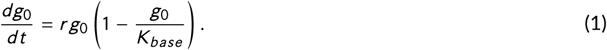

With the help of the secreted materials, cells can utilize new sources of nutrients,consequently, we assume that these leaky materials increase the carrying capacity of the population. Since the efficiency of the secreted material depends on its concentration, the cooperative term of the carrying capacity *K*_*coop*_ depends on the frequency of producers. We use a simple linear relationship between the frequency of producers and carrying capacity *K*_*coop*_ (Oliveira et al., 2014) given by

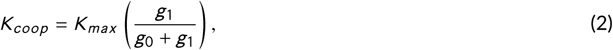

where *K*_*max*_ is a constant that describes the maximum carrying capacity which can be reached if all cells produce the secreted material. Thus, the presence of producers modifies the dynamics of the cheaters to

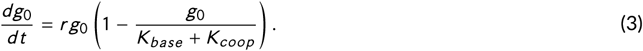

Producing leaky materials is costly, which means that the replication rate decreases or the death rate increases for the producer strain. Let *r*_1_ = *r* − *c* denote the net growth rate of producer cells, where *c* (*r* > *c* > 0) is the cost of producing the leaky material. We assume that producer and non-producer strains differ only in the production of the leaky material, that is, all cells are genetic variants of closely related strains and thus, they compete for the same limiting resources with the same efficiency. Consequently, the strength of competition among the individuals of both strains should be the same. This term is *r* /(*K*_*base*_ + *K*_*coop*_) for cheaters (see eq. 3), which should be the same for cooperator strains. In this way, the dynamical model of the cooperator-cheater system will be the following

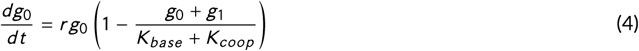

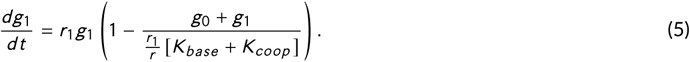

We note here that Oliveira et al. (2014) neglected the *r*_1_/*r* term in eq. (5) which cannot be supported by biological arguments and results in a structurally unstable model (for more details, see Discussion).

However, microbial populations are rarely well-mixed. Cells move in a limited and correlated manner even in liquid environments (Kümmerli et al., 2014; Sretenovic et al., 2017) and leaky materials diffuse with a limited speed or a fraction of them can bind to the surface of producers. A consequence of the nonrandom distribution of cells and the secreted material is that the relative fitness of cheaters and cooperators is density-dependent. Evidence that the relative fitness of cooperators can increase or decrease with density (see the Introduction) is supported by theoretical and experimental results. In this way, taking this effect into consideration we modify the availability of the secreted material in the function of density and trait-type in eqs. (4, 5). We introduce the Δ(*g*) function which describes the density-dependent private benefit of producers, where *g* = *g*_0_ + *g*_1_ is the total density of the population. The cooperators’ carrying capacity increases with Δ(*g*), while the cheaters’ carrying capacity decreases with this term. Thus we obtain the following dynamics for the cheaters and cooperators.

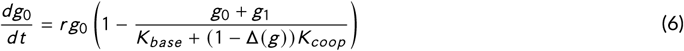

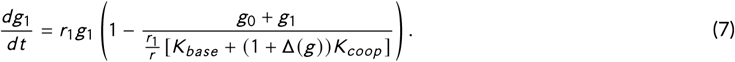

In line with experimental observations (Lindsay et al., 2018) we used a saturating function for Δ(*g*) if the private benefit of cooperators increases with density

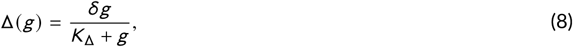

and a similar decreasing function if the benefit of cooperators decreases with density

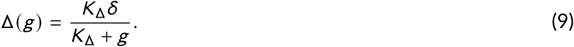

In both cases *δ* > 0 refers to the maximum intensity of the private benefit and *K*_Δ_ determines the density where this effect reaches half of the maximum value (Δ(*K*_Δ_) = *δ*/2) (Fig. 1).

**FIGURE 1.**
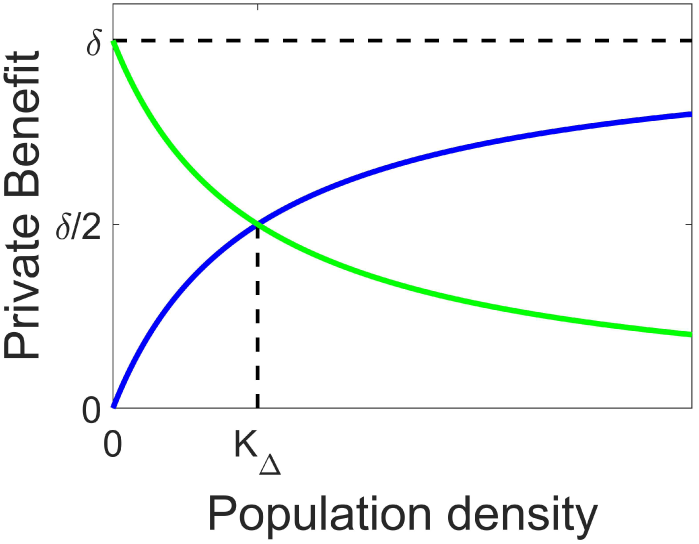
The private benefit of cooperators as a function of the population density. Private benefit either increases (blue) or decreases (green) with density. The functions are characterized by parameters *K*_Δ_ and *δ*.

### Population dynamics of traits with two different secretions

We consider the case when microbes can extract two different materials which determines four different genotypes: the cheater, producing neither material (denoted by [0,0]), the two types of specialist cooperators, producing one of the materials (denoted by [1,0] and [0,1]), and the generalist cooperator, producing both materials ([1,1]). With the same assumptions applied above, the dynamics of traits are described by

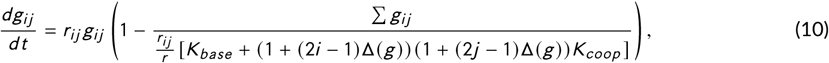

where *g*_*ij*_ represents the density and *r*_*ij*_ = *r* − [(*i* + *j*)]^*v*^*c* is the replication rate of genotype [*i, j*] and *g* = *g*_*ij*_ =∑ *g*_00_ + *g*_10_ + *g*_01_ + *g*_11_ equals to the total density.

We introduce a new parameter *v* > 0 to characterize the cost function of producing more different secretions. *v* refers to accelerating cost with the number of produced materials. *v* = 1 is selected in most cases, that is the cost is simply additive, but we also studied the effect of decelerating and accelerating costs as well. The two secretions can act in an additive, multiplicative, or synergistic (super-additive) ways on the carrying capacity (*K*_*coop*_). In the multiplicative case, both secretions can act only together. Then, coexisting specialist cooperators are in an interdependent mutualistic interaction in this case. Since we were interested in the possibility of interdependent mutualism between bacterial strains, we assumed that secretions act multiplicatively on *K*_*coop*_ (Oliveira et al., 2014). Thus we use the following multiplicative *K*_*coop*_ function

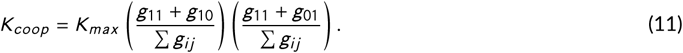

Similarly, we can define a dynamic model of strains producing any number of different leaky materials (see the Supplementary Material). Later we will study the behavior of 1-secretion, 2-secretions and 3-secretions models in detail.

### The structured population model

The above defined dynamical model describes the dynamics of cooperators and cheaters following a simple life cycle and living in an intensively mixed population. Density-dependent private benefit is the consequence of the limited diffusion of leaky materials and the limited motion of cells, therefore, some spatial structure is assumed in the background. This model reflects microbiomes living in aquatic or viscous but homogeneous habitats. However, numerous microbial communities follow a life cycle which can be modelled better by the so-called structured deme model (Wilson, 1975; Cremer et al., 2012; Oliveira et al., 2014; Chuang et al., 2009; Nadell et al., 2016). The life cycle has three phases in the model: i) *m* local populations are formed from the entire well-mixed population (invasion phase). The initial local population (group) size is determined by the Poisson distribution with mean *n* and individuals are selected randomly from the entire population and moved the local populations. ii) The local populations grow according to the dynamics defined above in eq. (10). iii) After the local populations reached their steady state (starvation phase), these subpopulations are merged to form a new population, with a new gene pool (sporulation and migration phase), as a starting point of a new lifecycle (Fig. 2).

**FIGURE 2.**
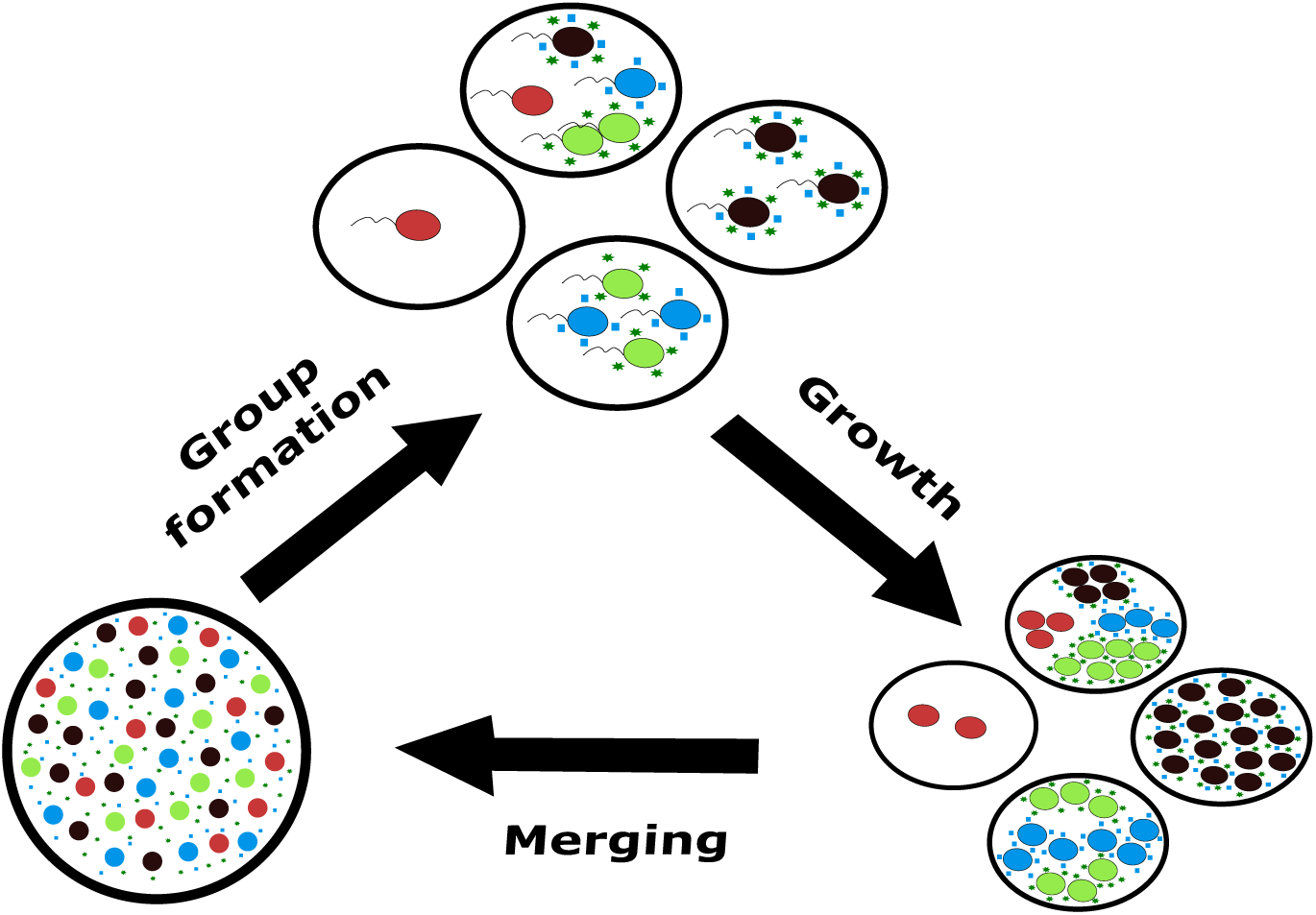
The structured deme model of the microbial lifecycle. The figure illustrates an example for the 2-secretions model. Each local population can contain either only specialists (green and blue circles), generalists (black circle) or cheater individuals (red circle) or any mixture of them in a group. Specialists can produce only certain secretions depending on their genotype (green stars and blue squares) while generalists produce both secretions. Each local population grows according to eq. (10) reaching a steady state, and then they merge, establishing new local populations randomly. This cycle repeats until the genotype frequencies reach a steady state for the whole community.

## 3 RESULTS

### Dynamical behavior of the local population

First, we studied the dynamical behavior of the 1-secretion model through analytical and numerical methods. Let us start the analysis with the case when *δ* = 0, where the system is well-mixed, and there is no private benefit. It is easy to show that cheaters always exclude cooperators from the population in this case, since *dg*_0_/*dt* > *dg*_1_/*dt* for every *g*_0_, *g*_1_ > 0. This result is not surprising since this is a dynamic model of the public goods game in a well-mixed environment. However, if *δ* > 0 then the system’s dynamical behavior becomes more complex. Assume that the cheater is the resident genotype and the cooperator is the invader. That is *g*_0_ → *K*_*base*_, while *g*_1_ → 0 and *K*_*coop*_ → 0. Thus

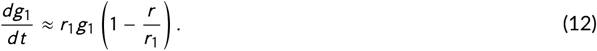

Since *r* > *r*_1_, *dg*_1_/*dt* < 0, this means that rare cooperators can never invade the population of cheaters. Similarly, rare cheaters cannot invade the population of cooperators if *dg*_0_/*dt* < 0 in the *g*_0_ → 0 limit. Since 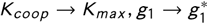, (where 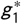 is the equilibrium density of the cooperators is the positive solution of 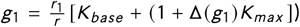 and 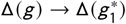, then by comparing eqs. (6) and (7) we have

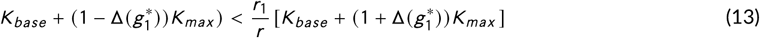

which supports *dg*_0_/*dt* < 0. After rearranging eq. (13), we find that cooperators are resistant against the invasion of rare cheaters if

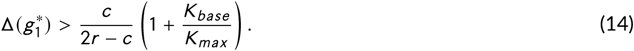

Consequently, if eq. (14) is valid then the monomorph cheater and monomorph cooperator populations are the alternative locally asymptotic states of the system. If eq. (14) is not true and Δ(*g*) is an increasing function then 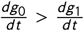 is always true, since *g*_0_ + *g*_1_ decreases due to the invasion of cheaters and Δ(*g*) also decreases. In this way, the cheater strategy is globally stable. The situation is different if Δ(*g*) is a decreasing function and eq. (14) is not true. Thus rare cheaters invade cooperators. However, cheaters’ invasion causes *g*_0_ + *g*_1_ to decrease and Δ(*g*) to increase. Consequently, the fitness of cheaters decreases while the fitness of cooperators increases. If there is a 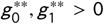 density pair where 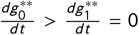 then this is a stable equilibrium point, in which cooperators and cheaters will be in a stable coexistence.

Based on the above results, we can give a sufficient condition when the cheater strategy invades and excludes the cooperator strategy. Since *δ* ≥ Δ(*g*), it follows from eq. (14) that if

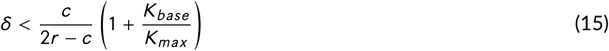

then cheaters win against cooperators. Without loss of generality, we might assume that *r* = 1. (A different *r* only rescales the time unit.) Assuming a low cost (*c* ≪ *r*) and a high cooperative carrying capacity compared to the basic (*K*_*base*_ ≪ *K*_*max*_) the above relation simplifies to *δ*/*c* < 1/2.

The qualitative analysis described above is confirmed and visualized by numerical simulations. Assuming that all parameters are fixed except *δ* and *c*, and assuming that *c* << *r* then eqs. (14) and (15) are determined only by *δ*/*c*, the *maximal relative private benefit of cooperation*. Fig. 3 presents the stable equilibria of eqs. (6) and (7) as a function of *δ*/*c*.

**FIGURE 3.**
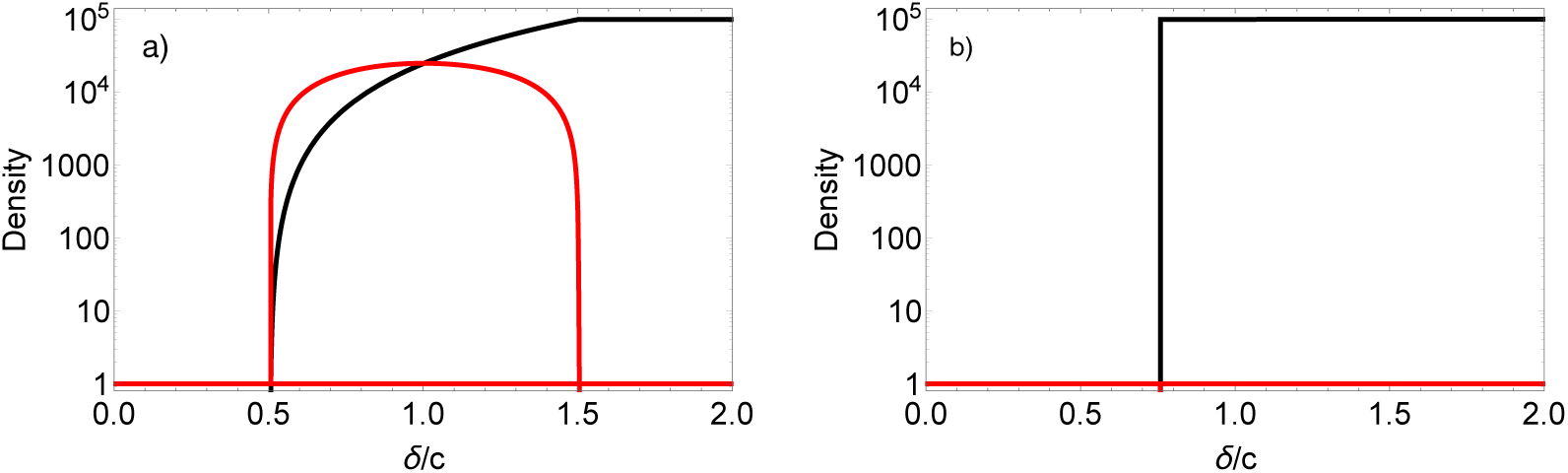
Stable fixed points of the 1-secretion model as a function of *δ*/*c*. Red lines denotes the equilibrium density of cheaters, black lines denotes the equilibrium density of cooperators. The cheater state is globally stable for *δ*/*c* below a critical level and locally stable above it (horizontal red line at density 1). a) Private benefit decreases with density. b) Private benefit increases with density. *c* = 0.01, *r* = 1, *K*_*base*_ = 1, *K*_*max*_ = 10^5^, *K*_Δ_ = *K*_*max*_ /2

If the private benefit decreases with density, then cheater strategy is the only fixed point of the dynamics if *δ*/*c* ≲ 1/2. There is a polymorphic stable equilibrium of cheaters and cooperators between 1/2 ≲ *δ*/*c* ≲ 3/2, while the cheater strategy becomes a locally stable state if *δ*/*c* ≳ 1/2. If *δ*/*c* ≲ 3/2 the system will be bi-stable with alternative monomorphic stable states, the dynamics converges either to the cooperative or to the cheater state (Fig. 3a). The situation is simpler if the private benefit increases with density. Then again the cheater strategy is the only stable state below a critical *δ*/*c*, and the system becomes bi-stable with the pure cooperative and pure cheater state above this *δ*/*c* level (Fig. 3b). Naturally, there is an unstable polymorphic fixed point between the alternative stable states, which is not depicted in the figure. For the analysis of the 2-secretions model, again it can be shown easily that cheaters win over cooperators if there is no private benefit (*δ* = 0). Using a similar argument from the 1-secretion model, we can give a sufficient condition for the competitive dominance of cheaters over generalist cooperators (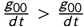, for every *g*_00_, *g*_11_ > 0). Knowing that Δ(*g*) ≤ *δ* cheaters are dominant over generalist cooperators if

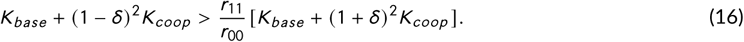

Assuming that *δ* ≪ 1 and thus *δ*^2^ ≪ *δ*, cheaters exclude the generalist cooperators if

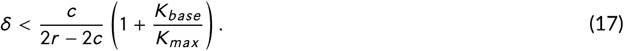

Similarly, cheater is dominant over the specialist cooperator [1,0] or [0,1] if

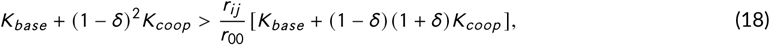

where *r*_*ij*_ = *r*_10_ or *r*_01_. We assume here that both specialist cooperators are present in the population, thus *K*_*coop*_ > 0. By neglecting the *δ*^2^ term as before the above relation leads to

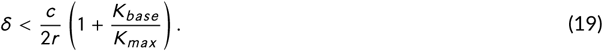

Since eq. (17) follows from eq. (19), eq. (19) guarantees the competitive superiority of cheaters over any cooperative strategy. Again, if *K*_*base*_ ≪ *K*_*max*_ and *r* = 1, then *δ*/*c* < 1/2 gives a sufficient condition for *δ*/*c* below which the cheater is the only stable state. The consequence of the above analysis is that if *δ*/*c* is greater than a critical value then the competitive superiority of the cheaters is not supported. We examine the local asymptotic stability of the single strategies in this case by studying their resistance against the rare invader strategies. Following the same way of thinking as in the 1-secretion model (see eq. 12) it can be shown that neither the rare generalist cooperator nor the pair of rare specialists can invade the population of cheaters.

Let us now consider the case when the generalist cooperator is the resident strategy. Specialist cooperators are effective only if they emerge together. The rare pair of [0,1], [1,0] strategies do not invade the resident generalist strategy if *dg*_10_/*dt* < 0 and *dg*_01_/*dt* < 0. Using eq. (10) in this special case, we observe that specialists cannot invade the generalist strategy if

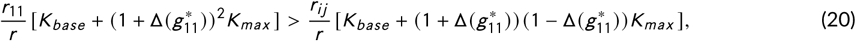

where *r*_*ij*_ refers to either *r*_10_ or *r*_01_ and 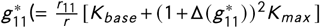 is the equilibrium density of the generalist strategy if it is alone in the population. Assuming again that *δ* ≪ 1, thus neglecting the Δ^2^ term and if

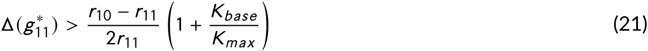

then the generalist cooperator strategy is stable against the invasion of rare specialists. Applying a very similar calculation we determine that the generalist cooperator is resistant against the invasion of the rare cheater strategy if

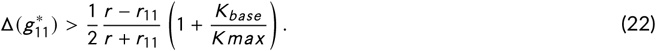

Consequently, if both eqs. (21) and (22) are valid, then the generalist cooperator strategy is stable against all possible rare invaders. Whether eq. (21) or (22) is the stricter condition depends on *v* and *c*. The linear case (*v* = 1) in eq. (22) follows from eq. (21), that is if the generalist strategy is resistant against the invasion of specialist cooperators then it is stable against cheaters as well.

Since we have already shown that rare cooperators never invade the cheater population, the remaining step is to study the stability of specialist cooperators against the invading generalist cooperators and cheaters. We know that the dynamics of strategy [0,1] and [1,0] are determined by the same equations, consequently 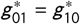 and thus *K*_*coop*_ = *K*_*max*_ /4 in the steady state, if the cheater and generalist cooperator strategies are missing from the system. After the same analysis as before, we conclude that specialist cooperator [0,1] is resistant against the invasion of the rare cheater strategy if

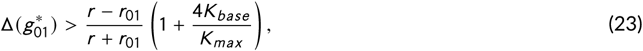

and resistant against the invasion of the rare generalist cooperator strategy if

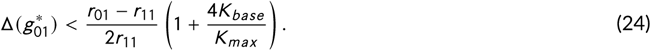

The same relations are valid for genotype [1,0], only the indices are changed. The above conditions can be satisfied simultaneously only if (*r* − *r*_01_/(*r* + *r*_01_) < (*r*_01_ − *r*_11_)/2*r*_11_, which can be true if *v* ≥ 1, that is when the cost is additive or accelerating.

The last option is that the specialist and generalist cooperators invade the population each others mutually. Generalist cooperators invade specialists when eq. (24) is not valid, and specialists invade generalists if eq. (21) is not true. These conditions can be satisfied if 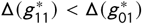. Since 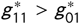 (except if *c* and *v* are unrealistically large), mutual invasion is possible if Δ(*g*) is a decreasing function. The consequence of mutual invasion is that generalist and specialist strategies could not exclude each other, but they coexist in this case.

The common nature of the above relations is that if all parameters are fixed except *c* and *δ* and it is assumed that *c* ≪ *r* then *δ*/*c* is the only parameter which determines the dynamics of the system. To demonstrate this, let us consider the eq. (22). After some rearrangement we have 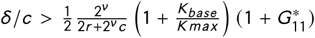, where 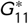 is either 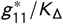 or 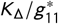 depending on whether the private benefit decreases or increases with density. Since *c* ≪ *r* the right hand side of the relation is approximately independent of *c*. The relation is determined only by *δ*/*c*.

Besides the qualitative analysis presented above, we performed numerical simulations to reveal the behavior of the system in a more comprehensive way and to show how the parameters affect the dynamics. The most important results are summarized in Fig. 4 and Fig. 5. We selected a basic parameter set and showed the stable states of the dynamics as a function of *δ*/*c* with this parameter set (Fig. 4 b and Fig. 5 b). In line with the mathematical analysis, the cheater strategy is the only stable state if *δ*/*c* is below a critical level. Above this level, the cheater strategy becomes only locally stable (a rare alternative strategy cannot invade) and alternative stable states emerge as well. When the private benefit decreases with density, first the cheater and the specialist cooperators will be in coexistence. We note that this stable state remains hidden in the local analysis since rare specialists never invades the resident cheater strategy. As *δ*/*c* increases further cheater strategy is dismissed by the generalist cooperator in the polymorphic equilibrium, and increasing *δ*/*c* even further results in that the generalist cooperator and the cheater will be the two alternative stable states of the dynamics (Fig. 4).

**FIGURE 4.**
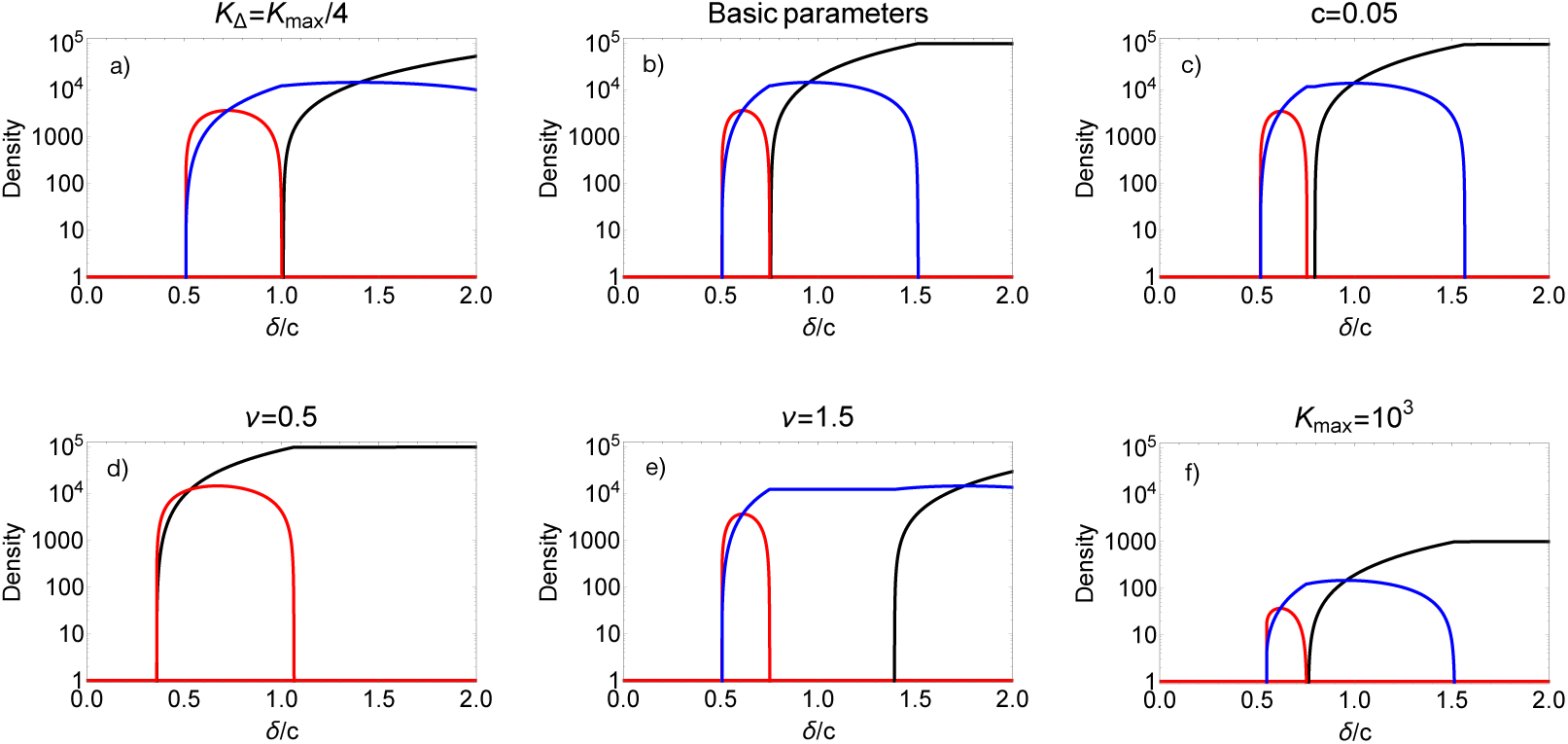
The stable states of the 2-secretions model as a function of maximal relative private benefit *δ*/*c*, when the private benefit decreases with density. Genotype [0,0] is denoted by red, [1,0] and [0,1] are denoted by blue, and [1,1] is denoted by black color. a) Stable densities at a smaller *K*_Δ_ (*K*_Δ_ = *K*_*max*_ /4). b) The stable densities at the basic parameters (*r* = 1, *c* = 0.01, *v* = 1, *K*_*base*_ = 1, *K*_*max*_ = 10^5^, *K*_Δ_ = *K*_*max*_ /2). c) Stable densities at a higher cost (*c* = 0.05). d) Stable densities when *v* is smaller than one (*v* = 0.5). e) Stable densities when *v* is bigger than one (*v* = 1.5). f) Stable densities at smaller *K*_*max*_ (*K*_*max*_ = 10^3^).

**FIGURE 5.**
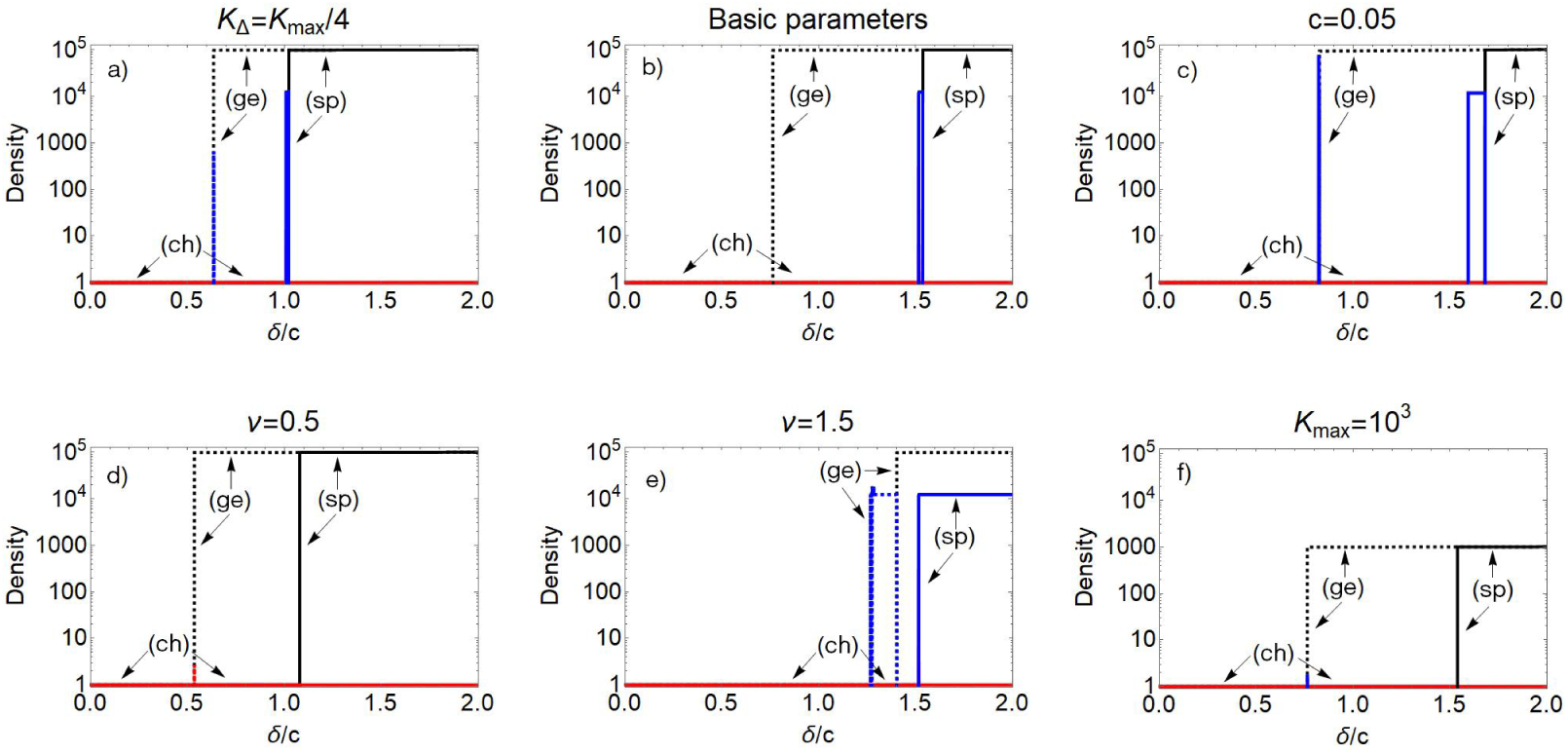
The stable states of the 2-secretions model as a function of maximal relative private benefit, *δ*/*c* when the private benefit increases with density. Genotype [0,0] is denoted by red, [1,0] and [0,1] are denoted by blue and [1,1] is denoted by a black color. Arrow (ch) indicates the bifurcation diagram when cheaters are dominant initially (*g*_00_ (0) = 1, *g*_10_ (0) = *g*_01_ (0) = *g*_11_ (0) = 0.1). In this case, cheaters always win the selection (horizontal red line). Arrow (ge) and dashed line indicate the bifurcation diagram when generalist is common initially (*g*_00_ (0) = 0.1, *g*_10_ (0) = *g*_01_ (0) = 0.1, *g*_11_ (0) = 10^4^). Arrow (sp) indicates the steady states when specialist cooperators are common initially (*g*_00_ (0) = *g* 11(0) = 0.1, *g*_10_ (0) = *g* 01(0) = 10^4^). There can be an interval of *δ*/*c* where specialists (blue) are resistant against the invasion of other strategies. Below that interval the cheaters and above that the generalist cooperators excludes the specialists. b) The stable densities at the basic parameters (*r* = 1, *c* = 0.01, *v* = 1, *K*_*base*_ = 1, *K*_*max*_ = 10^5^, *K*_Δ_ = *K*_*max*_ /2.) a) Stable densities at a smaller *K*_Δ_ (*K*_Δ_ = *K*_*max*_ /4). c) Stable densities at a higher *c* (*c* = 0.05). d) Stable densities when *v* is smaller than one (*v* = 0.5). e) Stable densities when *v* is bigger than one (*v* = 1.5). f) Stable densities at smaller *K*_*max*_ (*K*_*max*_ = 10^3^).

Again in harmony with the local mathematical analysis, the situation is totally different if the private benefit increases with density. The system remains bistable, that is, the equilibrium states depend on the initial densities, but there is no coexistence of different strategies. In Fig. 5, we depict the stable states starting the dynamics from three qualitatively different initial densities which correspond to either cheater, the generalist cooperator, or the specialist cooperators being the dominant strategies initially and the others are rare. As Fig. 5b shows if the cheater strategy is the dominant initially, then always the cheater wins the selection (line ch on Fig. 5). If generalist is the dominant initially then above a critical *δ*/*c* this strategy will be the winner, while below cheater wins (ge lines on Fig. 5). Last, in the case when specialists are dominant initially then cheater strategy wins for a wide range of *δ*/*c*, there is a narrow range of *δ*/*c* where specialists are the winner of the selection, and generalist strategy excludes the competitors for higher *δ*/*c* (sp lines on Fig. 5).

The effect of parameters on the dynamical behavior of the 2-secretions system is presented in the other panels of Fig. 4 and Fig. 5. We depict the same bifurcation diagrams, however *K*_Δ_, *c, v* and *K*_*max*_ were modified separately. As we have shown analytically, increasing the parameter *c* modifies the dynamical behavior of the system only very moderately at the small *c* limit (Fig. 4 c and Fig. 5 c). Similarly, changing the *K*_*max*_ value causes only minor changes in the behavior, still the *K*_*max*_ » *K*_*base*_ limit remains valid (Fig. 4 f and Fig. 5 f). As our mathematical analysis predicts, the generalist cooperator strategy is supported if *v* < 1 and suppressed if *v* > 1 (Fig. 4d,e and Fig. 5d,e). The effect of changing *K*_Δ_ is presented in (Fig. 4a and Fig. 5a). If the private benefit decreases with density, then decreasing *K*_Δ_ causes a faster loss of private benefits with density, this generalist strategy can be invaded by specialists strategies more easily (see eq. 21). Therefore, specialist strategies are present in stable coexistence with generalist strategies in a wider *δ*/*c* interval at higher *K*_Δ_ (compare Fig. 4a and b). In contrast, if the private benefit increases with density, then decreasing *K*_Δ_ increases the private benefit faster at smaller densities, and thus the generalist strategy is more successful against the invasion of the cheater strategy.

Our analysis also included the 3-secretions model. Since analytical derivation becomes even more complex for three traits and thus for eight genotypes, and the results remain qualitatively the same, we focus only on the numerical simulations. The stable states follow a similar pattern as in 2-secretions model. For convenience we introduce the following notations: Genotype [1,1,1], the generalist cooperator is denoted by *Co*3, genotypes [1,1,0], [1,0,1] and [0,1,1] are denoted jointly by *Co*2, and [1,0,0], [0,1,0], [0,0,1] by *Co*1 and [0,0,0], the cheater strategy is denoted by *Ch*.

If the private benefit decreases with density then the cheater strategy is the only stable state if *δ*/*c* ≲ 1/2. As *δ*/*c* increases further, the previously globally stable cheater strategy becomes only locally stable, and then, the coexistence of the *Ch* and *Co*1 emerges as an alternative stable state. As *δ*/*c* increases further,the *Ch* strategy is excluded from this stable polymorphic state, and the *Co*1 and *Co*2 strategies will coexist. For even larger *δ*/*c* values, *Co*1 is excluded by the *Co*3 and *Co*2 strategies which are in coexistence. Further increase of *δ*/*c* lead to the exclusion of the *Co*2 and the generalist cooperator (*Co*3) and cheater (*Ch*) strategies are the two alternative stable states (Fig. S1a). Similar to the 2-secretions model, the coexistence of strategies is not possible if the private benefit increases with population density. Below a critical level of *δ*/*c* the cheater strategy is the only stable state, while above this value one of the cooperator strategies (typically the generalist cooperator) becomes an alternative stable state (Fig. S1b).

### 3.1 Dynamical behavior in the structured population

In the applied structured population model it is common that strategies that increase the average fitness of the local population, can spread and even fixate in the whole population even though these can be competitively inferior to other strategies locally (Smith, 1964; Wilson, 1975; Chuang et al., 2009). In our model system, the cooperative genotypes are probably supported by the population structure. For the 1-secretion model the intuitive explanation is simple: consider a case when the cheater excludes the cooperator (*δ*/*c* < 1/2) in a local population. This will happen in every local population where there is at least one cheater initially. In those local populations which contain only cooperators initially, populations grow to a much higher population density (*K*_*max*_ ≫ *K*_*base*_) than in those “infected” by the cheaters. Consequently, the local cooperative populations send a lot of cooperators to the next generation, balancing or even overriding the weakness of cooperators in competition with cheaters locally. However, here we studied a more complex situation where besides the complete cheaters, the specialist and generalist cooperators are in competition with each other where the private benefit of the producers is density dependent.

To compare the dynamical behavior of the local population to the structured population, we used the same parameter set for the simulations as before. Further, we selected the *δ*/*c* values to scan the qualitatively different regimes of the dynamics. The total population is divided into *m* = 10^3^ local populations and the average number of founder individuals, *n* varies from 2 to 50 through the simulations (low and high heterogeneity). Initial densities are the same for all strategies during the simulations, however, the random selection of individuals into the local populations can form many different initial configurations and the local populations always start to grow from a different composition of genotypes.

According to the detailed numerical simulations in the 1-secretion model, cooperators exclude cheaters or are in a stable coexistence with them if a few individuals form a local population initially (roughly *n* < 10), while the equilibria are similar to those that follow from local population dynamics if there are numerous founder individuals (*n* = 50) (Fig. S2). The population structure clearly supports cooperation.

As we have shown above, there are qualitatively different regions in dynamical behavior of 2-secretions models. Thus, to receive a comprehensive picture about the dynamical behavior of the structured population, we selected *δ*/*c* = 0.3, 0.6, 1, and 2, when the private benefit decreases (see Fig. 4) and *δ*/*c*=0.5, 1, and 1.5, when the private benefit increases with density (see Fig. 5).

Fig. 6 summarizes the dynamical behavior of the structured populations. One of the most unambiguous characteristics of the selection dynamics in a structured population is that specialist cooperators will be the winners of the selection or a member of a coexisting community even under wider circumstances than in unstructured populations. Interestingly, specialist cooperators exclude the alternative genotypes even for a low maximal relative benefit when *n* is at the intermediate level. A very low *n* favours the generalist cooperators, while the cheater strategy excludes the others at a high *n* as in the local populations (Fig. 6 a,b). The explanation is the following: although generalist cooperators are competitively inferior to cheaters and specialist cooperators at a small *δ*/*c*, there is some probability that an initial local population is formed only by one genotype at very low *n* (*n* < 4). If only the generalist cooperators inhabit a local population initially then this population reaches a very high density (nearly *K*_*max*_) at the end of the growth phase, thus transferring a high number of generalists to the next generation. Naturally, it is highly probable that local populations will be founded by specialist cooperators and cheaters initially, however, specialist cooperators have a significantly smaller carrying capacity (nearly *K*_*max*_ /4) than the population of generalists, not to mention that the cheaters’ carrying capacity is *K*_*base*_. As a consequence, generalist cooperators exclude their competitors. As *n* increases, the probability of having a founder population with a single strategy decreases. This is detrimental to the generalist cooperators since they are competitively inferior to the other strategies while the specialist cooperators suffer less. Furthermore, higher *n* increases the probability that interdependent mutualistic couples be formed initially. Consequently, specialist cooperators win the competition for an intermediate *n*. As *n* becomes large (*n* ≥ 50) almost every local population contains at least one cheater strategy initially, which excludes the cooperators in the local population. Thus, the cheaters win as in the well-mixed local population.

**FIGURE 6.**
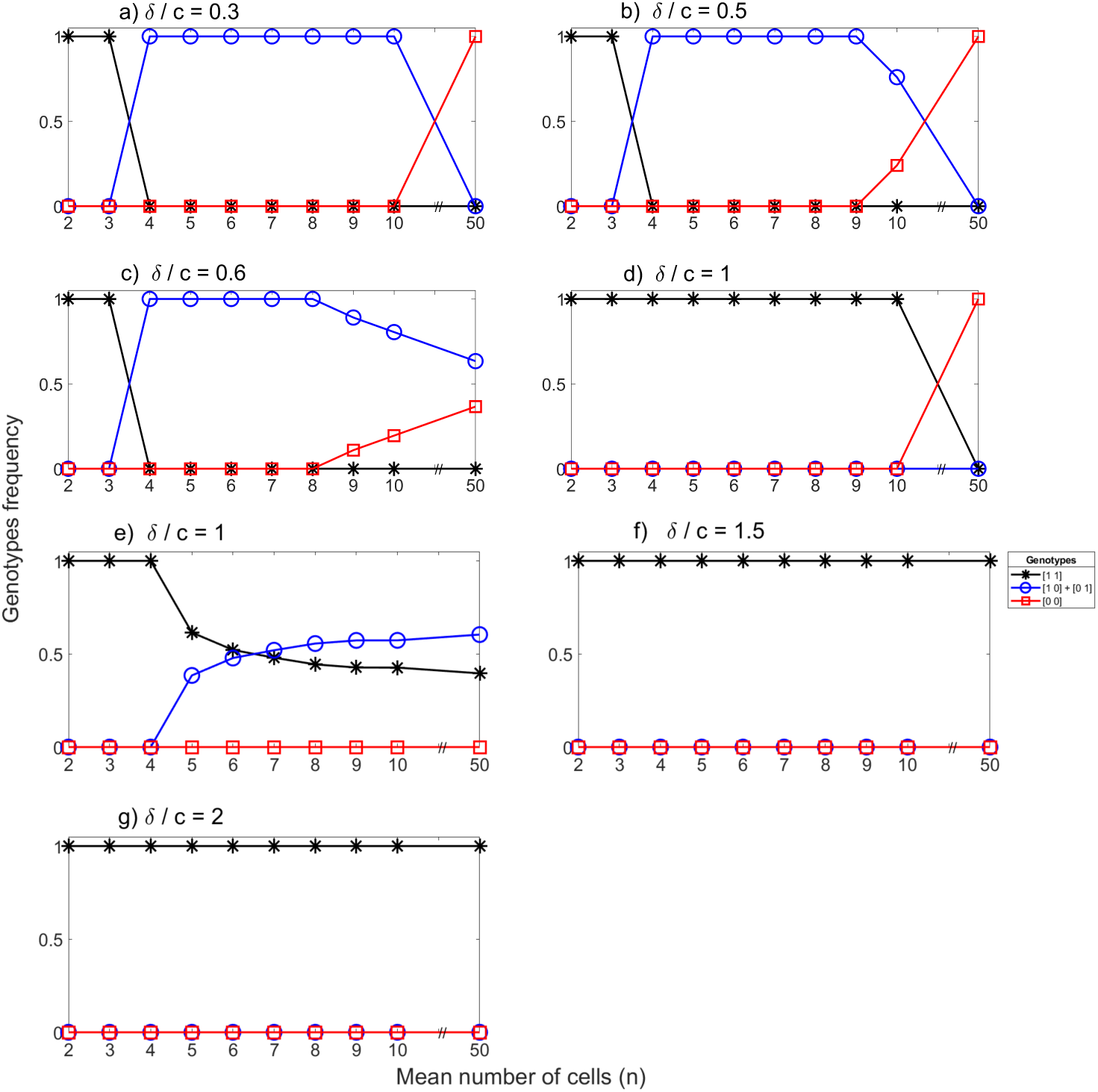
Selection among genotypes in differently mixing populations at characteristically different *δ*/*c*. Left column: the private benefit decreases with density. Right column: the private benefit increases with density. *δ*/*c* = 0.3, 0.6, 1, 2 in Figure (a),(c), (e) and (g), *δ*/*c* = 0.5, 1, 1.5 in Figure (b), (d), (f), respectively. The other parameters are the same as in Fig. 4 and Fig. 5.

Similarly to the single population model, specialist cooperators can coexist with either cheaters (Fig. 6c) or generalist cooperators (Fig. 6e) when the private benefit decreases with density and *δ*/*c* is not too small or not too high. According to the above argument, very low *n* is favorable for the generalist cooperators, while an intermediate level of *n* decreases the success of cheaters in competition with specialist cooperators (Fig. 6c). Population structure does not change selection in populations at high *δ*/*c* (Fig. 6g), that is the generalist cooperator wins the competition. When the private benefit increases with density and the *δ*/*c* ratio is above a critical level, then the generalist cooperators are the typical winners of the selection as in the single population model (Fig. 6d,f).

We analyzed how the dynamical behavior of the structured population depends on the other parameters of the model. Similar to the local population, decreasing *K*_Δ_ enhances the parameter space where specialist cooperators are favored (Fig. 7a,b). Likewise, the accelerating cost of producing multiple materials, that is if *u* > 1, favors the specialist cooperators (Fig. 7c, d). Naturally, changes of these parameters in the opposite directions weaken the success of specialists (not shown).

**FIGURE 7.**
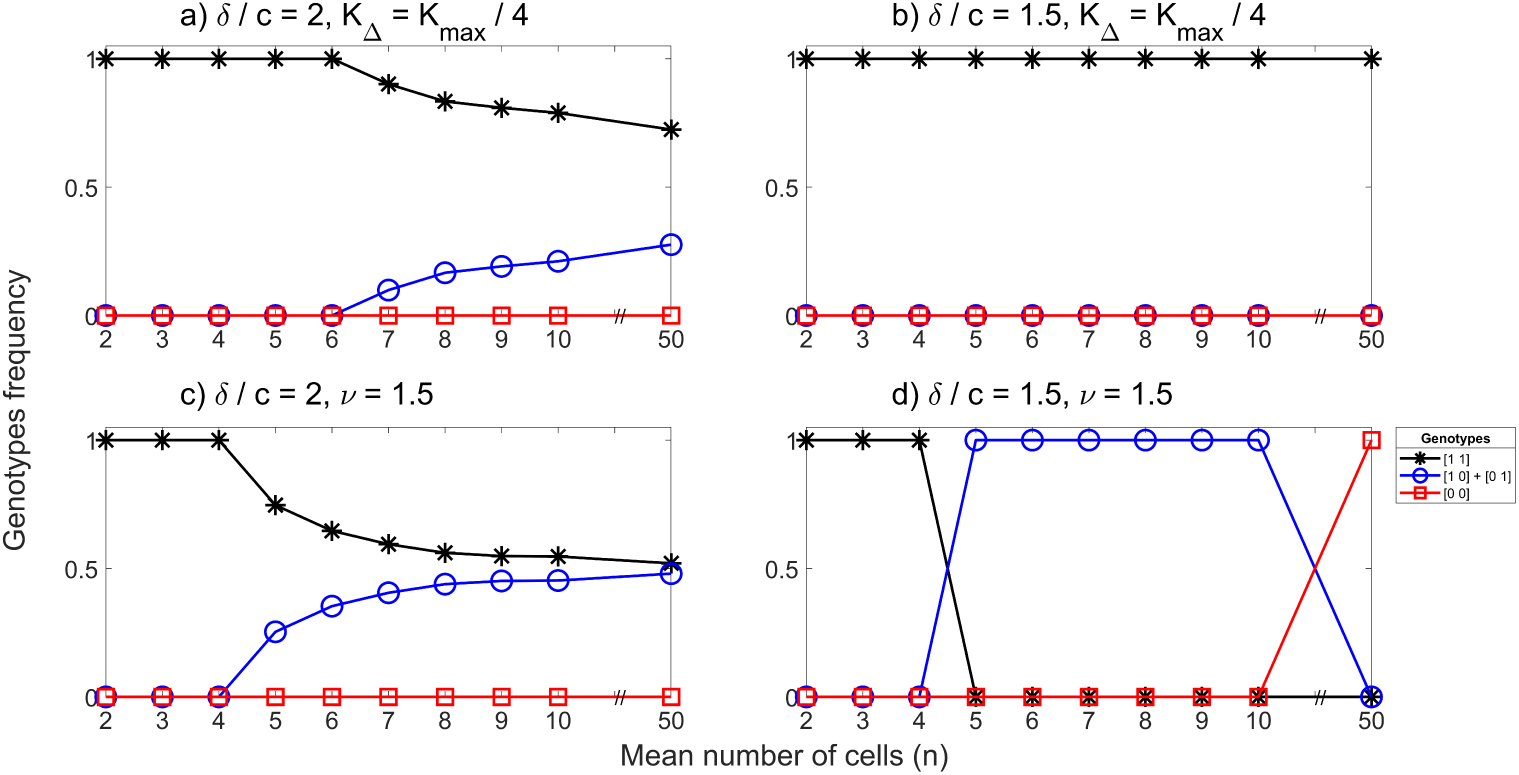
The effect of model parameters on selection. a), c) private benefit decreases with density, b), d) private benefit increases with density. a) A smaller *K*_Δ_ supports the interdependent mutualists if the private benefit decreases with density, c), d) an accelerating production cost (*v* > 1) favor interdependent mutualism. Changing the parameters in the opposite direction favors the generalist producer. Other parameters are the same as in Fig 6.

Knowing that the dynamical behavior of the local population for the 3-secretions model is similar to the 2-secretions model, it is not surprising that this qualitative similarity continues in the structured population. Similarly to the 2-secretions model we found that the *Ch* strategy wins the competition only if the relative private benefit of cooperation is small and the mean number of founder cells is high (Fig. S3), while *Co*3 always wins the competition if there are only two or three founder cells in the local population (Fig. S3). Further, if the relative private benefit is high, then *Co*3 will be the winner of the selection, independently of the number of founder cells (Fig. S3 g, h). *Co*3 is competitively inferior to any other strategy if *δ*/*c* is small or not big (Fig. S1 and Fig. S4), so this strategy is excluded in every local population where there are individuals from the other strategies initially. This happens more and more frequently as *n* increases. On the contrary, *Co*2 cooperators are successful when there are only a few founder individuals locally and *δ*/*c* is small or moderate (Fig. S3a,b,c). In these cases, there is a high probability that two different genotypes of *Co*2 are placed together in the same local population. They mutually help each other; thus this local population can reach a high population density. Although *Ch* or any member of *Co*1 excludes the mutualistic *Co*2 pairs (see Fig. S1 and Fig. S4), the chance of these competitors being together with the *Co*2 pairs in the founder populations is low if *n* is not too large. Increasing the number of founders further makes co-residence of *Ch* and *Co*1 with *Co*2 more probable initially. Consequently, the relative benefit of *Co*_2_ decreases. In parallel, the complete co-residence of the interdependent *Co*1 community occurs with higher probability in local populations, increasing the relative benefit of this strategy. Thus *Co*1 will be the winner of the selection or will be dominant in the whole population at a medium *n* (Fig. S3 a,b, c). As in the 2-secretions model, different cooperative strategies are in coexistence only if the private benefit decreases with density, *δ*/*c* is roughly 0.5 to 1.5 and *n* is not too small to prevent the emergence of cooperative coalitions initially (Fig. S3 c, e). According to the numerical simulations (not shown), the parameter dependence of the results remains the same as in the 2-secretions model: smaller *K*_Δ_ and *v* > 1 are favorable for the specialist cooperators, while *v* < 1 increases the parameter space where the generalist win the competition.

## 4 DISCUSSION

We studied a model of microbial mutualism where the competing cells can produce different types of leaky materials. The production of leaky materials was modeled to be costly but producing them serves as a private benefit for the producer cell, and this private benefit depends on population density. Further, we considered a situation when all of the different leaky materials are necessary to cause a positive effect and the specialist producers are completely interdependent on each other. Different analyses with competition between cell types, either the one’s non-producing leaky materials to the types which produce all different secretions were done, in an intensively mixed population with simple life cycle and in a structured population with a more detailed life cycle.

Naturally, if the private benefit is missing or weak compared to the production cost, then the situation is identical to the public goods game and then the cheater strategy is the only stable state in the intensively mixed population. However, as we showed with mathematical and numerical methods, mutually dependent cooperative strains are present in a steady state in a well mixed population for a broad range of conditions, if the private benefit decreases with density (Fig. 4, Fig S1a). Depending on the maximal relative private benefit (*δ*/*c*), the mutually dependent cooperative strains coexist either with more or less generalist cooperative strains (Fig. 4 and Fig. S1 a). These mutualist strains are either exploited by strains producing a lower number of different materials including the cheater strain, or these strains exploit the more generalist strain in the coexisting community. Fig. 8 demonstrates the interaction network of mutually dependent genotypes ([1,0] and [0,1]) coexisting either with the cheater ([0,0]) or with the generalist ([1,1]) genotype.

**FIGURE 8.**
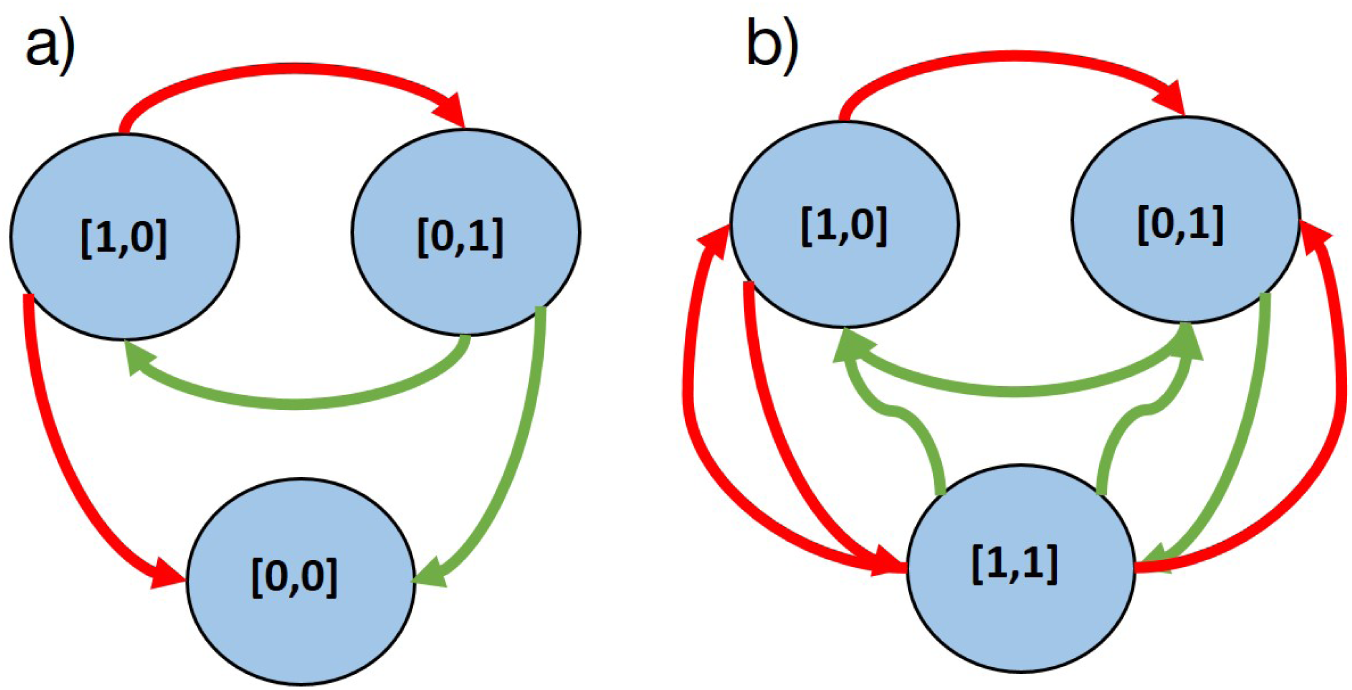
Interaction of mutually dependent cooperative strains with the cheater or the generalist cooperator in the 2-secretions model. Specialist cooperators [1,0] and [0,1] produce one of the two materials (red and green). a) The cheater genotype ([0,0]) exploits the cooperators, b) The generalist genotype ([1,1]) is exploited by the specialist cooperators.

In those cases, when the private benefit increases with density (and *δ*/*c* is above a critical level) the population has two equilibrium states. One of them is the cheater strategy and the other is typically the generalist cooperator strategy. Mutually dependent cooperative strains win the competition only under very specific circumstances (Fig. 5 and Fig. S1).

In agreement with the results of previous work (Oliveira et al., 2014; Estrela et al., 2016), if producing more secretions is more costly, that is if there is a tradeoff between the synthesis of the secretions, then the generalist strategy is suppressed and interdependent mutualism is supported. Contrary, if producing more secretions is less costly, that is if the synthesis of different secretions is present in multiple genotypes then the generalist strategy is more supported, and specialists are suppressed (Fig. 4, Fig. 5, Fig. 7).

The speed of change of density dependence when density is small is determined by *K*_Δ_ in our model. A quicker decrease (smaller *K*_Δ_) of density dependence of the private benefit helps the interdependent mutualists to coexist with the generalist (Fig. 4a,b and Fig. 7a). The situation is the opposite if the private benefit increases with density: faster saturation to the maximum private benefit increases the success of the generalists and suppresses the specialists (Fig. 5a,b and Fig. 7b). The intuitive explanation is the following: specialists have a competitive benefit over the generalists at lower densities since their density independent growth rate is larger than the generalist’s growth rate (*r*_10_ = *r*_01_ > *r*_11_), but the generalists can have a fitness benefit over specialists at higher densities because they can reach a higher equilibrium density. The relative dominance of these opposite effects is affected by the density dependence of the private benefit. If it is strong at lower densities and negligible at high densities, this supports the specialists, if it is significant at high densities this promotes the generalist strategy.

In similar work, Estrela et al. (Estrela et al., 2016) studied a model with a private benefit of secretions, where cells could produce at most two different secretions. Besides these similarities, there are significant differences between their work and our present model. They studied density independent private benefit, and, drove by the secretion of antibiotic degradation molecules, they assumed that these molecules decrease the death rate of the cells. Despite these differences in the models, they obtained results very similar to ours: mutual interdependency is favored at an intermediate level of private benefit, and the coexistence of different strains is present for a broad range of conditions. Hence, interdependent mutualism and coexistence of different strains are maintained by this private benefit under different biological circumstances. Our results are less comparable with the results of Oliviera et al. (2014), most importantly because no private benefit was included in their model. Furthermore, they used a population dynamical model in which all genotypes coexist in a neutrally stable equilibria in the local populations, which makes the model structurally unstable (Arnold, 1988).

Theoretical and experimental studies pointed out earlier that cooperators are supported in the spatially structured populations (Nowak, 2006; Chuang et al., 2009). However, these studies considered only the selection of a cooperative and a cheater strategy, different common goods and the mutual dependence of specialized cooperators, in other words, the presence of partial cheaters was not possible. One of the few exceptions is the work by Oliveira et al. (2014) who they focused on the possibility of mutualistic interactions between specialized cooperators in a structured population. They have found that microbial mutualism is possible only if the number of founder individuals is not extremely large (*n* = 4 − 8) and the cost of producing the leaky material is high (*c* > 0.1). This result led them to the conclusion that interdependent mutualism among the partners competing for the same niche should be rare in microbial communities. We used in our model the same population structure as Oliveira et al (2014), but our conclusion is significantly different because density dependent private benefit is taken into account. Comparing the dynamical behavior in the structured model with the well-mixed model (see e.g. Fig. 4, Fig. 5 and Fig. 6), it is apparent that dynamics in the local population are strongly retained in the structured population, although the population structure further increases the possibility of microbial cooperation. Naturally, if *n* is small then the generalist genotype will be the winner of the competition, but specialist mutualists win or coexist with other genotypes for a larger number of founder cells under very different circumstances (Fig. 6 Fig. S3). Further, the mutually dependent cooperators are selected in the structured population even in those cases when the cheater strategy is the winner of the competition in the well-mixed population. Our results suggest that interdependent mutualism should not be rare in real microbial communities, especially in those in which their private benefit decreases with density. The selection of interdependent cooperation and coexistence of traits with different levels of cooperation, in general, is in harmony with the Black Queen Hypothesis which explains the loss of essential leaky functions of microbes as an adaptive process (Morris et al., 2014; Morris, 2015). The consequence of the adaptive loss of the production of leaky materials is that the effectiveness of communities with specialist cooperators is sub-optimal since maximal population density can be attained if generalist cooperators dominate the community (Fig. 4).

The above conclusion is modulated by the fact that we studied a worst case scenario from the point of view of cross-feeding, since we assumed that all secretions are needed for the enhanced carrying capacity. Further, we assumed that genotypes use the same niche, in this way, competition is maximal between the different strategies. Ruling out these assumptions, the possibility of coexistence of different genotypes increases and we hypothesize that cross-feeding, the complex networks of more and less specialized cooperators, and complete cheaters will be even more widespread in these communities including those where private benefit increases with density. These questions could be explored in a further study.

In summary, the character of density dependence of the private benefit determines essentially how easily interdependent mutualism can evolve. If the private benefit decreases with density, then interdependent mutualist genotypes typically coexist with cooperative strains while if the private benefit increases with density then interdependent mutualisms are rare. These general conclusions are valid in intensively mixing populations with a simple life cycle and in a more complex life cycle.

## Supplementary Material

### The general model with any number of secretions

Let us assume that strains can produce *n* ≥ 1 number of different secretions. Let us denote a genotype of a strain with [*i*_1_, *i*_2_, …*i*_*n*_], where *i*_*k*_ ∈ {0, 1} denotes whether the actual material is secreted (*i*_*k*_ = 1) or is not secreted (*i*_*k*_ = 0) by the strain. Thus the general dynamical system is

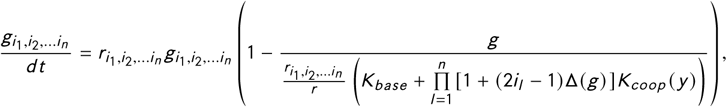

where 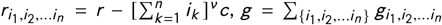 is the total density of strains (summation for {*i*_1_, *i*_2_, …*i*_*n*_}, means all combinations of index values),

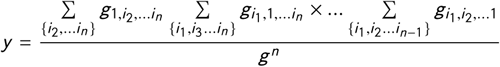

is the collective concentrations of secretions. *K*_*coop*_ (*y*) = *K*_*max*_ *S* (*y*), where *S* (*y*) is an arbitrary (linear or non-linear) function of *y*.

### 1-secretion and 3-secretions models

Here are the figures of 1-secretion and 3-secretions models, which we refer in the main text.

**FIGURE S1.**
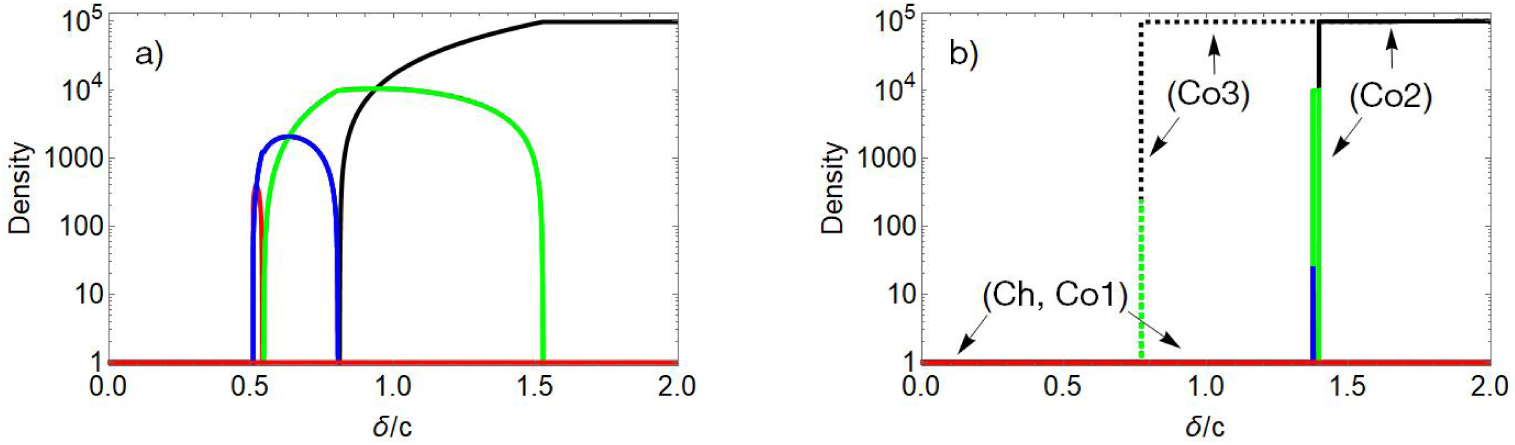
The stable states of the 3-secretions model in function of maximal relative private benefit *δ*/*c*. Genotype [0,0,0] is denoted by red, [1,0,0], [0,1,0] and [0,0,1] by blue, [1,1,0], [1,0,1] and [0,1,1] by green, [1,1,1] is denoted by black. a) stable states occur when private benefit decreases with density. b) Stable states occur when private benefit increases with density. Arrows (*Ch, Co*1) indicate the stable state when the cheater or *Co*1 strategies are common initially (*g*_*ij k*_ = 0.1, (*i, j, k*)*c* {0, 1}, except *g*_000_ (0) = 1, or *g*_*ij k*_ (0) = 0.001, (*i, j, k*)*c* {0, 1}, except *g*_110_ = *g*_101_ = *g*_011_ = 10^3^). Arrow (*Co*3) and dashed lines indicate the stable states when generalists are common (*g*_*ij k*_ (0) = 0.001, (*i, j, k*)*c* {0, 1}, except *g*_111_ (0) = 10^3^) and arrow (*Co*2) denotes the steady states when [1,1,0], [1,0,1], and [0,1,1] genotypes are common initially (*g*_*ij k*_ (0) = 0.001, (*i, j, k*)*c* {0, 1}, except *g*_110_ (0) = *g*_101_ (0) = *g*_011_ (0) = 10^3^). The parameters are the same as in the 2-secretions model.

**FIGURE S2.**
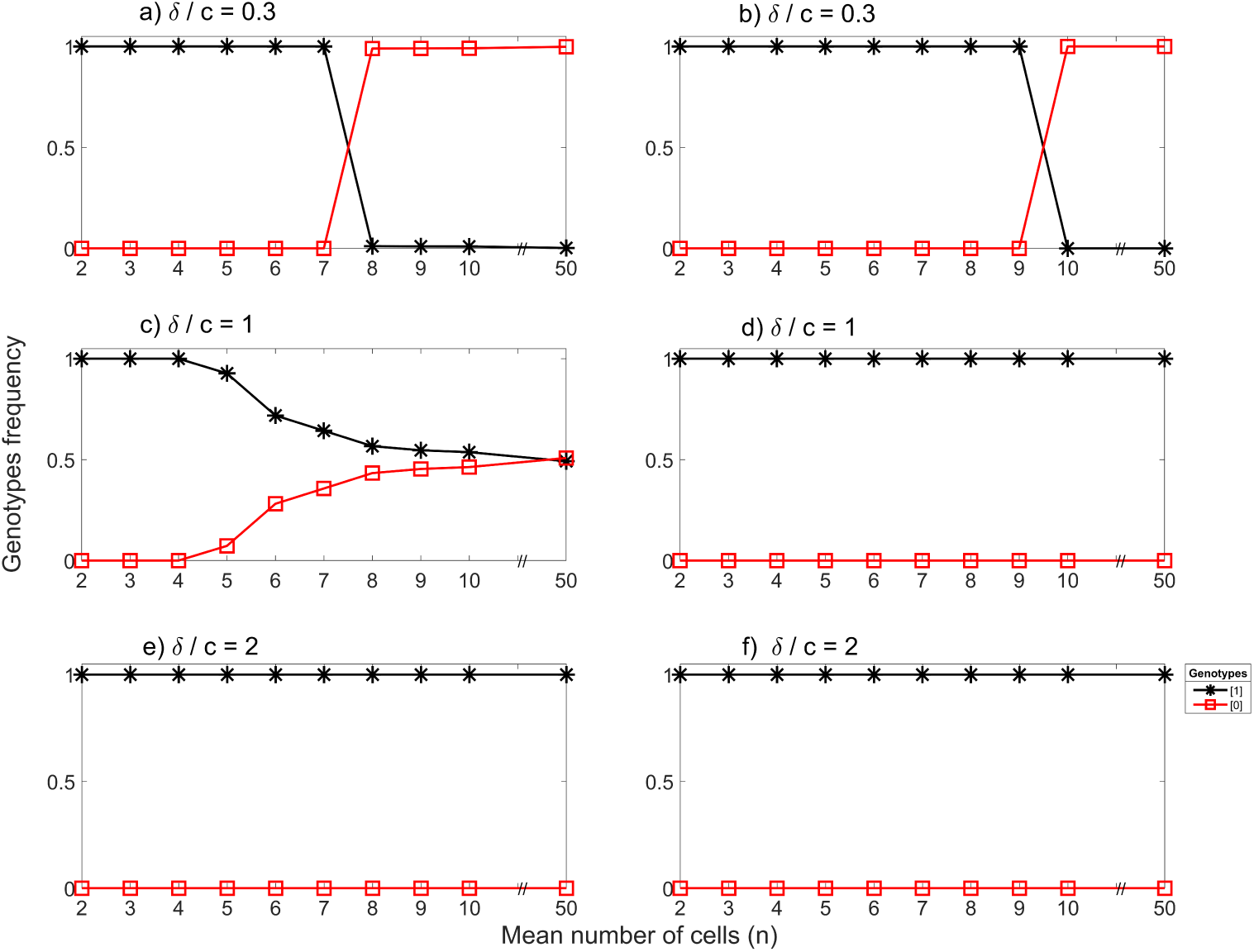
Selection between cheaters and cooperators in the structured 1-secretion model. (a), (c), (e) the private benefit decreases with density, (b), (d), (f) the private benefit increases with density. Black lines and stars denote the cooperator ([1]), red lines and squares the cheater ([0]). *δ*/*c* = 0.3, 1, 2 from top to bottom and other parameters are the same as before.

**FIGURE S3.**
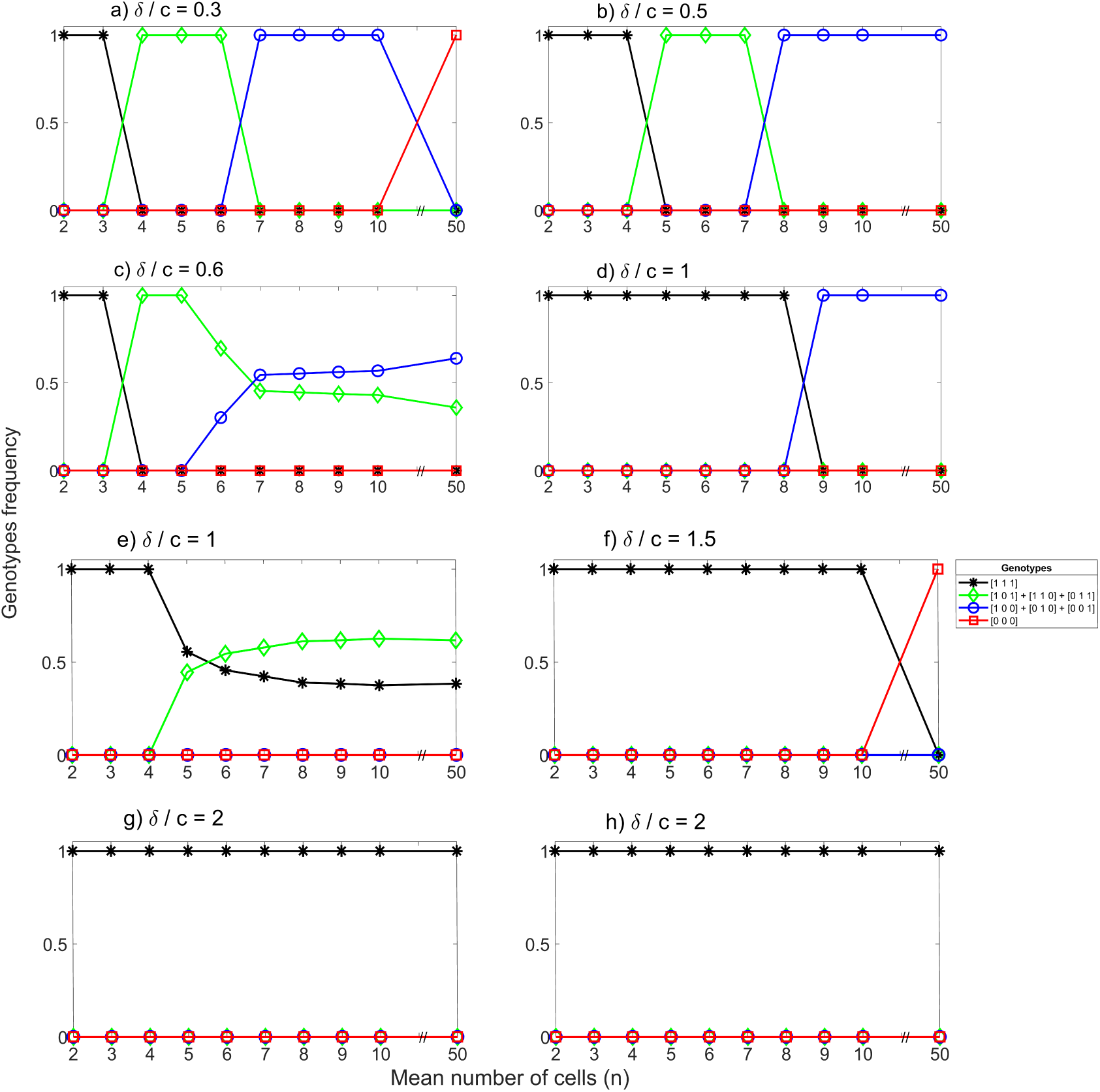
Selection among genotypes with 3 secretions in differently mixing populations. Left column: the private benefit decreases with density. Right column: the private benefit increases with density. *δ*/*c* = 0.3, 0.6, 1, 2 in Figure (a),(c), (e) and (g), *δ*/*c* = 0.5, 1, 1.5, 2 in Figure (b), (d), (f), (h), respectively. The other parameters are the same as in Fig.4 and Fig. 5.

**FIGURE S4.**
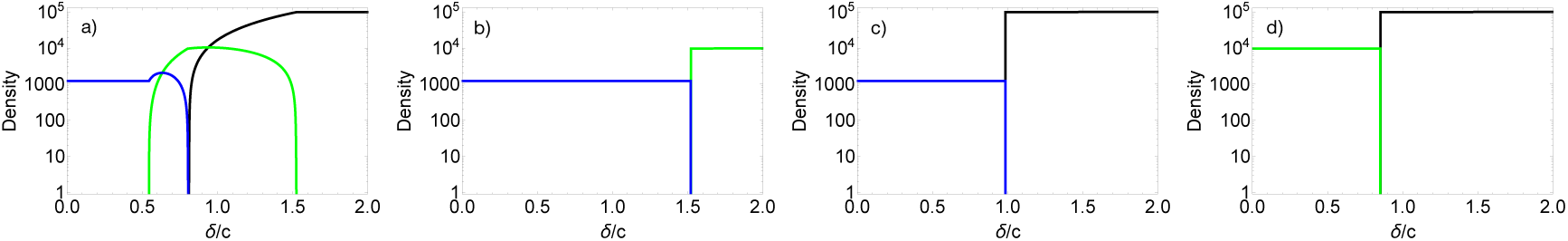
Competition between the cooperators if *Ch* is missing. a) The private benefit decreases with density. All cooperative strategies are in competition. Initial densities are 0.1 for all strategies. b), c), d) The private benefit increases with density and pairwise competition. b) Competition between *Co*1 and *Co*2. c) Competition between *Co*1 and *Co*3, d) Competition between *Co*2 and *Co*3. Initial density is 1 for the more generalist strategy and 0.1 for less generalist strategy in all cases while other strategies are not included. Color code: *Co*1 (blue), *Co*2 (green), *Co*3 (black).

## references

Archetti, M. and Scheuring, I. (2012) Game theory of public goods in one-shot social dilemmas without assortment. J Theor Biol, 9–20.

Archetti, M. (2016) Evolution of optimal Hill coefficients in nonlinear public goods games. J Theor Biol, 73–82.

Arnold, V. (1988) Geometrical Methods In The Theory Of Ordinary Differential Equations. Springer–Verlag.

Bardgett, R., Freeman, C. and Ostle, N. (2008) Microbial contributions to climate change through carbon cycle feedbacks. The ISME journal, 805–814, doi.org/10.1038/ismej.2008.58.

Birer, C., Tysklind, N., Zinger, L. and Duplais, C. (2017) Comparative analysis of dna extraction methods to study the body surface microbiota of insects: A case study with ant cuticular bacteria. Mol Ecol Resour., e34–e45 https://doi:10.1111/1755-0998.12688.

Bourne, D. G., Morrow, K. M. and Webster, N. S. (2016) Insights into the coral microbiome: underpinning the health and resilience of reef ecosystems. Annu. Rev. Microbiol., 317–340. https://doi:10.1146/annurev-micro-102215-095440.

Cavaliere, M., Feng, S., Soyer, O. and Jiménez, J. (2017) Cooperation in microbial communities and their biotechnological applications. Environ Microbiol., 2949–2963. https://doi:10.1111/1462-2920.13767.

Cho, I. and Blaser, M. J. (2012) The human microbiome: at the interface of health and disease. Nat. Rev. Genet., 260–270. https://doi:10.1038/nrg3182.

Chuang, J. S., Rivoire, O. and Leibler, S. (2009) Simpsons paradox in a synthetic microbial system. Science, 323, 272–275.

Cordero, O., Ventouras, L., DeLong, E. and Polz, M. (2012) Public good dynamics drive evolution of iron acquisition strategies in natural bacterioplankton populations. Proc Natl Acad Sci U S A, 20059–20064. https://doi:10.1073/pnas.1213344109.

Coyte, K., Schluter, J. and Foster, K. (2015) The ecology of the microbiome: Networks, competition, and stability. Science, 350, 663–666.

Cremer, J., Melbinger, A. and Frey, E. (2012) Growth dynamics and the evolution of cooperation in microbial populations. Scientific Reports, [281].https://doi.org/10.1038/srep00281.

Czárán, T. and Hoekstra, R. (2009) Microbial communication, cooperation and cheating: quorum sensing drives the evolution of cooperation in bacteria. PLoS One, https://doi:10.1371/journal.pone.0006655.

Dobay, A., Bagheri, H., Messina, A., Kümmerl, R. and Rankin, D. (2014) Interaction effects of cell diffusion, cell density and public goods properties on the evolution of cooperation in digital microbes. J Evol Biol., 1869–1877.

Drescher, K., Nadell, C. D., Stone, H. A., Wingreen, N. and Bassler, B. L. (2014) Solutions to the public goods dilemma in bacterial biofilms. Curr Biol., 50–55. https://doi:10.1016/j.cub.2013.10.030.

Estrela, S., Morris, J. and Kerr, B. (2016) Private benefits and metabolic conflicts shape the emergence of microbial interdependencies. Environ Microbiol, 1415–1427. https://doi:10.1111/1462-2920.13028.

Foster, K. (2004) Diminishing returns in social evolution: the not-so tragic commons. J. Evol. Biol., 1058–1072.

Foster, K. and Bell, T. (2012) Competition, not cooperation, dominates interactions among culturable microbial species. Curr Biol., 22, 1845–1850.

Gore, J., Youk, H. and van Oudenaarden, A. (2009) Snowdrift game dynamics and facultative cheating in yeast. Nature, 253–256. https://doi:10.1038/nature07921.

Harcombe, W. (2010) Novel cooperation experimentally evolved between species. Evolution, 2166–2172.https://doi:10.1111/j.1558-5646.2010.00959.x.

Harcombe, W., Riehl, W., Dukovski, I., Granger, B., Betts, A., Lang, A., Bonilla, G., Kar, A., Leiby, N., Mehta, P., Marx, C. and Segrè, D. (2014) Metabolic resource allocation in individual microbes determines ecosystem interactions and spatial dynamics. Cell Rep., 1104–1115. https://doi:10.1016/j.celrep.2014.03.070.

Helbing, D., Szolnoki, A. and Perc, M. and Szabó, G. (2010) Evolutionary establishment of moral and double moral standards through spatial interactions. PLoS Comput Biol, e1000758. https://doi:10.1371/journal.pcbi.1000758.

Hillman, E., Lu, H., Yao, T. and Nakatsu, C. (2017) Microbial ecology along the gastrointestinal tract. Microbes Environ., 300–313 https://doi:10.1264/jsme2.ME17017.

Jacoby, R., Peukert, M., Succurro, A., Koprivova, A. and S, K. (2017) The role of soil microorganisms in plant mineral nutrition-current knowledge and future directions. Front Plant Sci., 1617 https://doi:10.3389/fpls.2017.01617.

Kaeberlein, T., Lewis, K. and Epstein, S. (2002) Isolating “uncultivable” microorganisms in pure culture in a simulated natural environment. Science, 296, 1127–1129, https://doi10.1126/science.1070633.

Kokko, H. and López-Sepulcre, A. (2007) The ecogenetic link between demography and evolution: can we bridge the gap between theory and data? Ecol. Lett., 773–782.

Kümmerli, R., Schiessl, K., Waldvogel, T., McNeill, K. and Ackermann, M. (2014) Habitat structure and the evolution of diffusible siderophores in bacteria. Ecol Lett., 17(12), 1536–1544. https://doi:10.1111/ele.12371.

Lindsay, R., Pawlowska, B. and Gudelj, I. (2018) When increasing population density can promote the evolution of metabolic cooperation. ISME J, 849–859. https://doi.org/10.1038/s41396-017-0016-6.

Mattoso, T., Moreira, D. and Samuels, R. (2012) Symbiotic bacteria on the cuticle of the leaf-cutting ant acromyrmex subterraneus subterraneus protect workers from attack by entomopathogenic fungi. Biol Lett., 461–464. https://doi:10.1098/rsbl.2011.0963.

Mee, M., Collins, J., Church, G. and Wang, H. (2014) Syntrophic exchange in synthetic microbial communities. Proc Natl Acad Sci U S A, E2149–E2156. https://doi:10.1073/pnas.1405641111.

Morris, J. (2015) Black queen evolution: the role of leakiness in structuring microbial communities. Trends Genet., 31(8), 475–482. https://doi:10.1016/j.tig.2015.05.004.

Morris, J., Papoulis, S. and Lenski, R. (2014) Coexistence of evolving bacteria stabilized by a shared black queen function. Evolution, 68(10), 2960–2971. https://doi:10.1111/evo.12485.

Nadell, C. D., Drescher, K. and Foster, K. R. (2016) Spatial structure, cooperation and competition in biofilms. Nature Reviews Microbiology, 14, 589–600.

Nowak, M. A. (2006) Five rules for the evolution of cooperation. Science, 1560–1563. https://doi.org/10.1126/science.1133755.

Oliveira, N., Niehus, R. and Foster, K. (2014) Evolutionary limits to cooperation in microbial communities. Proc Natl Acad Sci, 17941–17946.

Pande, S., Merker, H. and Bohl, K. e. a. (2014) Fitness and stability of obligate cross-feeding interactions that emerge upon gene loss in bacteria. ISME J., 953–962. https://doi:10.1038/ismej.2013.211.

van der Ploeg, J. (2005) Regulation of bacteriocin production in streptococcus mutans by the quorum-sensing system required for development of genetic competence. J. Bacteriol., 3980–3989.

Rankin, D. (2007) Resolving the tragedy of the commons: the feedback between population density and intraspecific conflict. J. Evol. Biol., 173–180.

Rivière, A., Gagnon, M., Weckx, S., Roy, D. and De Vuyst, L. () Mutual cross-feeding interactions between bifidobacterium longum subsp. longum ncc2705 and eubacterium rectale atcc 33656 explain the bifidogenic and butyrogenic effects of arabinoxylan oligosaccharides. Appl. Environ. Microbiol.

Ross-Gillespie, A., Gardner, A., Buckling, A., West, S. and Griffin, A. (2009) Density dependence and cooperation: theory and a test with bacteria. Evolution., 2315–2325.

Sherwin, E., Bordenstein, S., Quinn, J. and Dinan, TG Cryan, J. (2019) Microbiota in social brain. Science, eaar2016 https://doi:10.1126/science.aar2016.

Smith, J. M. (1964) Group selection and kin selection. Nature, 201, 1145–1147.

Smith, N. W., Shorten, P. R., Altermann, E., Roy, N. C. and McNabb, W. C. (2019a) The classification and evolution of bacterial cross-feeding. Frontiers in Ecology and Evolution, 7, 153. URL: https://www.frontiersin.org/article/10.3389/fevo.2019.00153.

Smith, R., Doiron, A., Muzquiz, R. and et al. (2019b) The public and private benefit of an impure public good determines the sensitivity of bacteria to population collapse in a snowdrift game. Environ Microbiol., 21(11).

Sretenovic, S., Stojkovi, B., Dogsa, I., Kostanjšek, R., Poberaj, I. and Stopar, D. (2017) An early mechanical coupling of planktonic bacteria in dilute suspensions. Nat Commun., https://doi:10.1038/s41467-017-00295-z.

Stewart, E. (2012) Growing unculturable bacteria. J of Bacteriology, 194, 4151–4160.

Szabó, G. and Hauert, C. (2002) Phase transitions and volunteering in spatial public goods games. Phys Rev Lett, 118101.

Tarnita, C. (2017) The ecology and evolution of social behavior in microbes. Journal of Experimental Biology, 220, 18–24. https://doi:10.1242/jeb.145631.

Taylor, V. (2019) The microbiome and mental health: Hope or hype? J Psychiatry Neurosci., 219–222. https://doi:10.1503/jpn.190110.

Thompson, J. R., Rivera, H. E., Closek, C. J. and Medina, M. (2015) Microbes in the coral holobiont: partners through evolution, development, and ecological interactions. Front. Cell. Infect. Microbiol., 176. https://doi:10.3389/fcimb.2014.00176.

Valdes, A., Walter, J., Segal, E. and Spector, T. (2018) Role of the gut microbiota in nutrition and health. BMJ, k2179 https://doi:10.1136/bmj.k2179.

Wilson, D. S. (1975) A theory of group selection. Proceedings of the National Academy of Sciences, 72, 143–146.

Yurtsev, E., Chao, H., Datta, M., Artemova, T. and Gore, J. (2013) Bacterial cheating drives the population dynamics of cooperative antibiotic resistance plasmids. Mol Syst Biol, 683 https://doi:10.1038/msb.2013.39.

